# Cross-Tissue Concordance of Blood-Trained Protein EpiScores in Donor-Matched GTEx Tissues

**DOI:** 10.64898/2026.10.20.760574

**Authors:** Paulina Gonzalez Orozco, Fredrick Schumacher

## Abstract

**Background:** Protein EpiScores are DNA methylation (DNAm)-derived predictors of circulating protein levels that may provide accessible biomarkers of disease-related molecular variation. However, because these models were developed in blood, it remains unclear whether they preserve interindividual variation when applied to other tissues. We evaluated the cross-tissue concordance of blood-trained protein EpiScores in matched donors from the Genotype-Tissue Expression (GTEx) project.

**Methods:** We applied 109 published protein EpiScores to DNAm from whole blood and six matched non-blood tissues in 54 donors contributing 158 blood–tissue pairs. Fifteen scores had unusable variation, leaving 94 estimable scores per tissue and 564 comparisons. Positive concordance required Pearson r and Spearman ρ ≥0.50; strong concordance required both ≥0.70. Benjamini–Hochberg false-discovery-rate (FDR) correction was applied separately across all comparisons for each method. Leave-one-out analyses assessed donor influence, and partial Pearson correlations evaluated sensitivity to available covariates. Protein annotations, composites, and donor strata were examined descriptively.

**Results:** Ninety-seven of 564 comparisons (17.2%) met the positive-concordance criterion, and 73 of these (75.3%) met the leave-one-out stability criteria. Lung showed the broadest correspondence: median absolute Pearson r was 0.594, 56/94 scores were concordant, and 49 were stable. After covariate adjustment, median absolute partial r in lung was 0.522 and 68/94 scores had global q <0.05. Among initially concordant pairs, 48/56 in lung retained partial r ≥0.50 and q <0.05, compared with 3/10 in colon, 1/2 in kidney cortex, and 0/3 in prostate. Breast and ovary were too small for adjusted analyses. Exploratory donor-cohort differences did not survive FDR correction.

**Conclusions:** Blood-trained protein EpiScores showed broad correspondence in lung that persisted after adjustment for available covariates. Initially concordant findings in the other tissues eligible for adjustment were mostly covariate-sensitive. These results prioritize score–tissue combinations for replication and molecular validation; they do not establish that blood EpiScores measure local tissue protein abundance.

## Introduction

DNA methylation (DNAm) biomarkers can characterize variation in aging and disease-related processes. Early epigenetic clocks predicted chronological age across tissues [1], while subsequent models incorporated mortality-related information: GrimAge includes DNAm surrogates for smoking and selected plasma proteins [2], and PhenoAge estimates a mortality-related phenotype derived from clinical biomarkers [3]. These developments expanded the use of DNAm to study molecular signatures beyond chronological age.

Protein EpiScores estimate protein-associated signatures from weighted combinations of methylation-array measurements. Gadd et al. developed predictors for 953 plasma proteins and identified 109 EpiScores that met independent-test performance criteria, explaining approximately 1%–58% of variation in corresponding protein measurements after adjustment for known protein quantitative trait loci [4]. Application to 9,537 Generation Scotland participants identified associations with incident diseases over follow-up of up to 14 years [4]. These models enable investigation of protein-associated methylation signatures in cohorts with DNAm data, although they remain statistical proxies rather than direct measurements of protein abundance.

One proposed advantage of methylation-based protein proxies is their potential to capture sustained biological influences that a single circulating protein measurement may incompletely represent [4,5]. However, greater temporal stability cannot be assumed across predictors. A longitudinal comparison of five methylation-based C-reactive protein markers found that stability over approximately a decade varied by marker and was not consistently greater than that of measured CRP [6]. These findings support evaluating EpiScores as potentially complementary biomarkers whose properties must be established for each intended application.

Disease studies illustrate their potential relevance. Protein EpiScores have been associated with invasive breast cancer risk [7] and colorectal cancer survival [8]. In the latter study, adding selected EpiScores to clinical models improved prognostic discrimination in a sample of 136 patients, although independent validation remains necessary [8]. Using proteins measured by mass spectrometry, Robertson et al. also identified EpiScores associated with incident cardiovascular disease [5]. These applications motivate further investigation of the biological variation represented by protein-associated methylation signatures, including whether that variation is shared across tissues.

A central question is whether blood-trained EpiScores preserve between-donor variation when applied to DNAm from other tissues. Findings from epigenetic clocks show that cross-tissue comparability cannot be presumed: Apsley et al. observed substantial within-person differences and generally low correlations between blood-derived and oral-tissue estimates for many clocks [9]. Direct evaluation of protein EpiScores outside blood has also begun. Schmunk et al. examined reliability across matched blood and saliva samples, including blood-derived protein EpiScores, within a framework for developing a saliva-based inflammatory biomarker [10]. However, it remains unclear how individual protein EpiScores correspond between blood and multiple internal tissues within the same donors.

Here, we evaluated 109 blood-trained protein EpiScores in donor-matched blood and six non-blood tissues from the Genotype-Tissue Expression (GTEx) project. We quantified cross-tissue correspondence using Pearson and Spearman correlations, assessed sensitivity to individual donors and available covariates, and examined exploratory donor strata. Target-protein secretion annotations and functional-group composites provided descriptive biological context. Our objective was to characterize the extent and limits of cross-tissue EpiScore concordance and identify score–tissue combinations for further validation, while distinguishing correspondence between methylation -derived scores from prediction of local tissue protein abundance. This study assessed methylation-derived score correspondence; RNA-seq integration and cell-type deconvolution were outside its scope.

## Methods

### Study design and data source

We conducted a cross-sectional analysis of donor-matched DNA methylation (DNAm) data to evaluate blood–tissue correspondence of protein EpiScores. Publicly available processed DNAm data were obtained from GEO accession GSE213478, the multi-tissue GTEx resource described by Oliva et al. [11]. The source dataset comprises 1,000 samples from nine tissues and 424 donors. Our analysis included whole blood paired with transverse colon, lung, kidney cortex, prostate, breast mammary tissue, or ovary. Samples were matched separately for each blood–tissue comparison using GTEx donor identifiers. Donors could contribute to more than one tissue comparison.

### DNAm preparation and protein EpiScore generation

DNAm profiling and initial quality control were performed by the source investigators using the Illumina Infinium MethylationEPIC array, as described previously [11]. Processed DNAm β values were organized into tissue-specific matrices with CpGs as rows and samples as columns. Each of the seven tissue input matrices contained 754,119 CpG sites. The blood matrix included 54 individuals.

Protein EpiScores were generated on June 29, 2026, using the hosted MethylDetectR Shiny application (https://shiny.igc.ed.ac.uk/MethylDetectR/) [12]. Full processed matrices were uploaded together with corresponding sex and age metadata, without filtering to the platform’s optional CpG truncation list. The analysis included the 109 protein EpiScores developed by Gadd et al. [4], which estimate protein-associated methylation signatures from weighted combinations of DNAm β values. Models were applied without retraining in GTEx. The hosted MethylDetectR application version identifier was not recorded.

According to the MethylDetectR documentation, missing methylation values at CpG sites present in an input dataset are imputed using the site-specific mean across input individuals. Predictor CpGs absent entirely from the input are assigned their mean β values from the Generation Scotland reference data [12]. This substitution permits score calculation but does not recover donor-specific variation at absent sites. CpG availability and the extent of reference-value substitution were not quantified separately for each protein EpiScore.

Numerical score exports were harmonized across tissues for downstream analysis. Original predictor names were retained, and non-protein predictors and reference variables were excluded from protein-specific summaries. Donor age in years was obtained from the original GTEx metadata. Numerical ages were checked for consistency across tissues and agreement with recorded age ranges before use in covariate-adjusted and age-stratified analyses. CHIT-1 denotes the MethylDetectR score for chitotriosidase, encoded by CHIT1; original score identifiers are used consistently in the results and figures.

### Individual-score correlations and concordance criteria

Eligibility was assessed separately within each matched blood–tissue dataset and required usable variation in both blood and tissue scores. Scores with zero or otherwise unusable variation were retained in quality-control and supplementary outputs as non-estimable. Each eligible comparison used the available complete donor pairs; full-tissue analyses included 9–45 matched donors.

Pearson correlation quantified linear association, and Spearman correlation quantified monotonic association between paired blood and tissue EpiScores. Pearson 95% confidence intervals were calculated using Fisher’s z transformation. Two-sided p-values were adjusted using the Benjamini– Hochberg procedure separately for Pearson and Spearman correlations across all 564 estimable protein–tissue comparisons. A globally adjusted q-value <0.05 defined statistical significance for each method. Within-tissue adjusted q-values were retained as supplementary outputs.

Positive concordance required both Pearson r and Spearman ρ ≥0.50; strong positive concordance required both coefficients ≥0.70. Classifications used unrounded estimates. Negative coefficients were retained as inverse associations and did not meet these positive-concordance criteria. Magnitude-based classifications were evaluated separately from statistical significance and did not require a p-value or false-discovery-rate threshold.

Tissue summaries report median absolute correlations and their 25th and 75th percentiles (Q1 –Q3). These percentiles describe variation across the fixed set of estimable scores. Counts and percentages meeting concordance criteria used the number of estimable scores within each tissue as the denominator. Breast and ovary analyses were considered exploratory because of their small matched samples. The same correlation methods, concordance criteria, and global FDR correction were applied to breast and ovary; their exploratory designation did not alter the testing procedure.

Supplementary Tables S1–S6 list comparisons significant by either correlation method and distinguish significance in both methods from significance in Pearson or Spearman alone. These tables were filtered from the globally adjusted results without redefining the multiple-testing family. Complete results for all 109 scores per tissue, including non-estimable scores, are provided in Supplementary Data S1.

### Leave-one-out sensitivity analysis

Leave-one-out analyses assessed the influence of individual donors on blood–tissue correlations. For every estimable protein–tissue comparison, Pearson and Spearman correlations were recalculated after sequentially omitting one matched donor pair. We recorded the minimum and maximum coefficients and the maximum absolute change from each full-sample estimate.

Stability classification was evaluated among comparisons meeting the full-sample positive-concordance criterion. A comparison was classified as stable when both coefficients remained ≥0.50 after every omission and the maximum absolute change in each coefficient was <0.30. These criteria assessed sensitivity within the observed sample and were not interpreted as independent validation. Comparisons with discordant Pearson and Spearman estimates were examined descriptively. Influential donors were not excluded from the primary analysis; full-sample estimates were retained for reporting.

### Covariate-adjusted sensitivity analysis

Partial Pearson correlations were estimated by separately regressing blood and tissue EpiScores on the same selected donor covariates and correlating the residuals. Candidate covariates were age, sex, body mass index (BMI), donor cohort, race, and ethnicity. Sex was omitted for sex-specific tissues. Covariate inclusion depended on availability and variation rather than association significance. Categorical covariates required at least two observed levels with at least three observations per level.

Analyses required at least 15 complete donor pairs and n ≥ 5k + 5, where k denotes the effective covariate degrees of freedom, calculated as the rank of the covariate design matrix minus the intercept. Tests used n−k−2 degrees of freedom. Comparisons with constant inputs or unusable residual variation were excluded. Benjamini–Hochberg correction was applied across all 376 estimable partial-correlation tests. Because eligible covariate sets differed across tissues, adjusted estimates were interpreted as tissue-specific sensitivity analyses.

### Secretion annotations and functional-group assignments

Target-protein annotations were manually curated from the Human Protein Atlas, accessed in March 2026 [13], and linked to all 109 EpiScores using an exact-name mapping. Five mutually exclusive secretion categories were used: secreted to blood; locally or extracellularly secreted but not to blood; not secreted to blood; other or ambiguously secreted; and unclear or missing annotation. Family-level or ambiguous annotations were retained as ambiguous rather than assigned to a specific gene without supporting evidence. Evidence of plasma detection was recorded separately from secretion classification. The mapping and category-level results are provided in Supplementary Data S2. The Human Protein Atlas release used for annotation was not recorded; the retained mapping is provided to document the annotations applied.

Each predictor was also assigned to one primary functional group using a manually curated mapping. The ten groups were cell adhesion/vascular/extracellular matrix; chemokines/cytokines/growth factors; coagulation/platelet/hemostasis; development/signaling/unclear; hormone/endocrine/binding proteins; immune response/inflammatory defense; lectins/carbohydrate binding; metabolic/lysosomal/enzymatic; neural/axon guidance; and proteases/protease inhibitors.

Individual-score results were summarized within each secretion category and functional group. Summaries included counts and percentages meeting the positive-concordance, strong-concordance, and leave-one-out stability criteria. Denominators comprised the estimable scores in the corresponding tissue and category. Annotations provided descriptive biological context; these analyses were not pathway-enrichment tests.

### Donor-level composite analyses

Composite scores were constructed separately for secretion categories and functional groups within each matched blood–tissue dataset. Each constituent EpiScore was standardized separately in blood and tissue by subtracting its matched-sample mean and dividing by its matched-sample standard deviation. Available standardized values were then averaged within each donor and category to obtain paired blood and tissue composites.

Scores with zero or otherwise unusable standard deviations could not be standardized and did not contribute to composite means. Missing standardized values were excluded from the corresponding donor-level average. The number of contributing scores was recorded, and category sizes were reported to identify composites based on few predictors. Pearson and Spearman correlations were calculated between paired donor-level composites. These analyses were descriptive and did not constitute formal tests of differences between categories.

For an additional descriptive analysis, functional-group composites were centered within each tissue comparison and pooled across the six non-blood tissues. Donors contributing to multiple tissue comparisons were represented more than once. Pooled correlations were therefore reported without inferential p-values or confidence intervals that would treat these observations as independent. A separate across-tissue summary calculated median absolute composite correlations among tissues with at least 10 matched donors. Tissue-specific estimates remained the primary composite results.

### Exploratory stratified analyses

Individual-score correlations were examined within strata defined by age, sex, donor cohort, and recorded ethnicity. Age strata used the median age of matched donors within each tissue: 49 years for lung and prostate, 53 years for transverse colon, 57.5 years for kidney cortex, 55 years for breast mammary tissue, and 56 years for ovary. Younger strata included donors at or below the tissue-specific median; older strata included donors above it.

The donor cohort distinguished organ procurement organization donors from postmortem donors. Sex and ethnicity strata used the available donor metadata, with ethnicity retained as the original recorded codes. Stratum sample sizes represented distinct matched donors rather than repeated donor–score records.

Within-stratum correlations required at least 10 complete donor pairs and usable score variation. Median absolute Pearson correlations summarized association magnitude irrespective of direction. Smaller strata were retained in the sample-size audit but did not contribute correlation estimates. Age, sex, and ethnicity comparisons were descriptive. Absolute sample correlations tend to be larger in smaller samples even without stronger underlying associations; differences in median absolute correlations across unequal strata were therefore not interpreted as evidence of biological differences.

### Exploratory comparisons between donor cohorts

Following inspection of the descriptive stratified results, formal exploratory comparisons of organ-donor and postmortem Pearson correlations were conducted in lung and transverse colon. Each comparison required at least 15 complete donors in both cohorts, nonconstant blood and tissue scores, and finite Fisher-transformed correlations.

For each eligible EpiScore, the difference between independent cohort correlations was evaluated using Fisher’s r-to-z transformation. Two-sided p-values were calculated from the standard normal distribution. Raw correlation differences were reported as organ donor minus postmortem, and confidence intervals for differences in Fisher-transformed correlations were retained on the Fisher-z scale. Differences in Spearman correlations were summarized descriptively.

Benjamini–Hochberg adjustment was applied jointly across all 188 eligible cohort-comparison tests, comprising 94 EpiScores in each tissue. An adjusted p-value <0.05 defined statistical support for a cohort difference. Failure to meet this threshold was not interpreted as evidence of equivalence between cohorts.

### Software and reproducibility

Downstream analyses were performed in R version 4.4.2 on a high-performance computing cluster. The reproducible workflow includes input validation, donor matching, Pearson and Spearman correlations, multiple-testing adjustment, leave-one-out sensitivity analysis, stratified and partial correlations, donor-level composites, and reconciliation checks.

Publication summaries and revised figures were generated from these outputs using Python, pandas, NumPy, SciPy, and Matplotlib. Dependency versions and input-file hashes are recorded in the accompanying build manifest. Hosted MethylDetectR score generation was a separate upstream step. The accompanying scripts regenerate downstream results from the required score exports and metadata; they do not automatically reproduce the hosted scoring step.

## Results

### Matched samples and EpiScore eligibility

Matched blood and non-blood samples were available from 54 unique GTEx donors, comprising 45 blood–transverse colon pairs, 37 blood–lung pairs, 33 blood–prostate pairs, 24 blood–kidney cortex pairs, 10 blood–breast mammary tissue pairs, and 9 blood–ovary pairs. Together, these represented 158 blood–tissue pairs, with some donors contributing to multiple tissue comparisons. No duplicate donor–tissue–score records were identified.

All 109 protein EpiScores were available in each tissue. The same 15 scores had zero or otherwise unusable variation in blood or the corresponding tissue across the six comparisons, leaving 94 estimable scores per tissue and 564 protein–tissue comparisons. Non-estimable scores were retained in the complete supplementary results to distinguish unavailable correlation estimates from weak associations.

### Lung showed the broadest observed blood–tissue concordance

Lung had the highest median absolute Pearson correlation, 0.594 (Q1–Q3, 0.450–0.734), and median absolute Spearman correlation, 0.555 (Q1–Q3, 0.404–0.722). Median absolute Pearson correlations were 0.318 in transverse colon, 0.286 in kidney cortex, 0.264 in ovary, 0.218 in breast mammary tissue, and 0.092 in prostate (Table 1). These summaries describe association magnitude; the heatmaps retain the direction of individual correlations (Figure 1A; Supplementary Figure S1). Comparisons between tissues were descriptive because matched donor sets and sample sizes differed; no formal tests of between-tissue differences were performed.

**Figure 1A.**
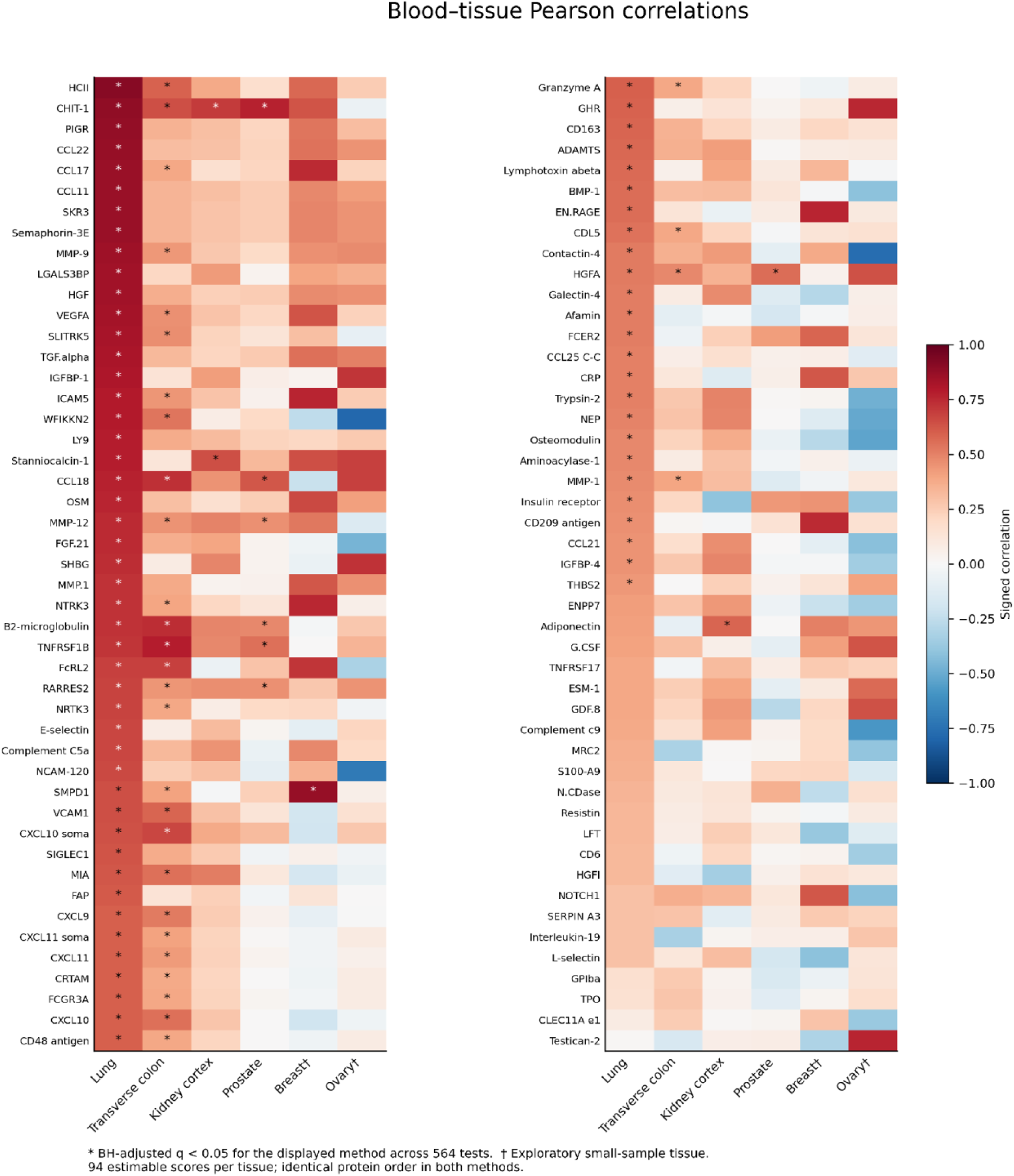
Individual-score Pearson correlations. Signed Pearson correlations for 94 estimable EpiScores, ordered by lung Pearson r. Asterisks indicate global BH-adjusted q <0.05 for Pearson across 564 tests. † Breast and ovary are exploratory. The matching Spearman display is Supplementary Figure S1.

**Table 1.** Blood–tissue concordance of protein EpiScores.

| Tissue | Matched donors, n | Median \|Pearson r\| (Q1–Q3) | Median \|Spearman ρ\| (Q1–Q3) | Both ≥0.50 n (%) | Both ≥0.70 n (%) | Both significant n (%) |
| --- | --- | --- | --- | --- | --- | --- |
| Lung | 37 | 0.594 (0.450–0.734) | 0.555 (0.404–0.722) | 56 (59.6) | 26 (27.7) | 67 (71.3) |
| Transverse colon | 45 | 0.318 (0.136–0.393) | 0.328 (0.158–0.404) | 10 (10.6) | 3 (3.2) | 26 (27.7) |
| Kidney cortex | 24 | 0.286 (0.173–0.403) | 0.260 (0.150–0.348) | 2 (2.1) | 0 (0.0) | 1 (1.1) |
| Prostate | 33 | 0.092 (0.036–0.216) | 0.110 (0.061–0.172) | 3 (3.2) | 1 (1.1) | 6 (6.4) |
| Breast mammary tissue† | 10 | 0.218 (0.068–0.483) | 0.248 (0.142–0.452) | 15 (16.0) | 4 (4.3) | 1 (1.1) |
| Ovary† | 9 | 0.264 (0.106–0.452) | 0.283 (0.117–0.446) | 11 (11.7) | 3 (3.2) | 0 (0.0) |

Correlations compare paired blood and tissue EpiScores. Of 109 protein EpiScores assessed per tissue, 94 were estimable and 15 had unusable variance. Percentages use 94 as the denominator. Median absolute correlations and Q1–Q3 describe the distribution across protein scores; Q1–Q3 are not confidence limits. Both ≥0.50 and both ≥0.70 require positive Pearson r and Spearman ρ to meet the specified threshold using unrounded values; the latter is a subset of the former. Both significant requires global Benjamini–Hochberg q < 0.05 for each method, with adjustment separately across 564 tests per method, regardless of correlation magnitude or direction. † Exploratory small-sample tissue. Donors may contribute to multiple tissues.

Overall, 97 of 564 comparisons (17.2%) met the positive-concordance criterion of both Pearson r and Spearman ρ ≥0.50. Lung accounted for 56 qualifying comparisons, corresponding to 59.6% of its 94 estimable scores. The corresponding counts were 15 in breast mammary tissue (16.0%), 11 in ovary (11.7%), 10 in transverse colon (10.6%), three in prostate (3.2%), and two in kidney cortex (2.1%) (Table 1; Figure 1B).

**Figure 1B.**
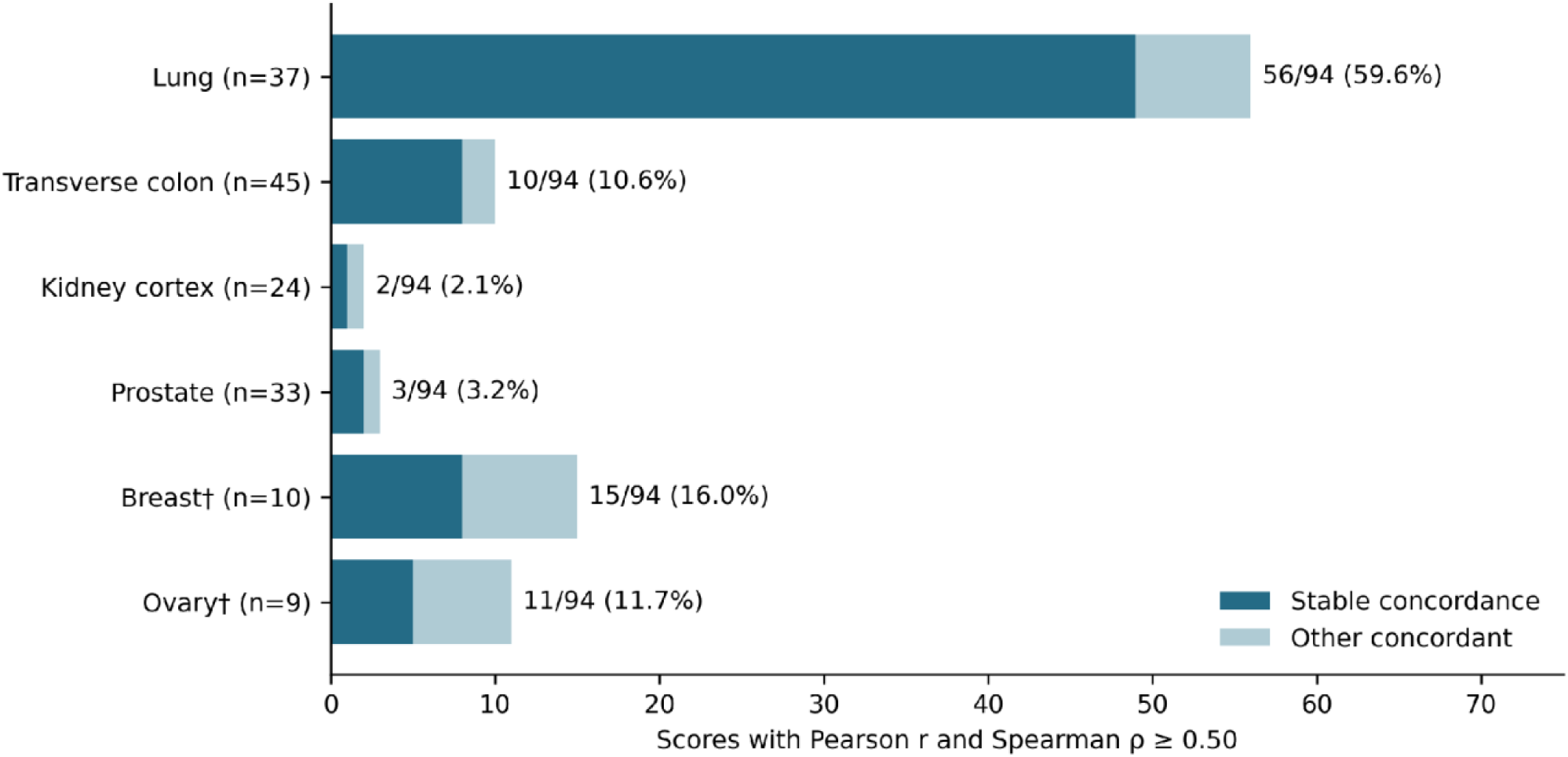
Concordance and donor sensitivity. Bars show scores with both r and ρ ≥0.50. The darker segment meets the combined leave-one-out stability rule: both coefficients ≥0.50 after every donor omission and maximum absolute changes <0.30. Percentages use 94 estimable scores.

Thirty-seven comparisons (6.6%) met the stronger criterion of both coefficients ≥0.70: 26 in lung, four in breast mammary tissue, three each in transverse colon and ovary, and one in prostate. No kidney cortex comparison met this criterion. These classifications describe positive correlation magnitude and consistency between methods, independently of statistical significance.

After global Benjamini–Hochberg adjustment, 134 comparisons were significant in at least one method: 101 in both methods, 13 in Pearson only, and 20 in Spearman only. The numbers significant in either method were 74 in lung, 42 in transverse colon, eight in prostate, four each in kidney cortex and breast mammary tissue, and two in ovary. Of these, 67 lung, 26 colon, six prostate, one kidney, and one breast comparison were significant in both methods; neither ovary comparison was significant in both. Supplementary Tables S1–S6 provide the tissue-specific lists, signed correlation estimates, adjusted q-values, concordance classifications, and stability results. Breast and ovary findings remained exploratory because of their small matched samples.

### Most concordant comparisons met the leave-one-out stability criteria

Of the 97 full-sample concordant comparisons, 73 (75.3%) met the combined stability criteria: both coefficients remained ≥0.50 after every single-donor omission, and the maximum absolute change in each coefficient was <0.30. The remaining 24 comparisons failed at least one criterion.

Lung accounted for 49 stable comparisons, followed by transverse colon and breast mammary tissue with eight each, ovary with five, prostate with two, and kidney cortex with one (Figure 1B). Thus, 49 of the 56 initially concordant lung comparisons (87.5%) retained concordance under the specified leave-one-out criteria. The 13 stable comparisons in breast and ovary remained exploratory despite meeting these operational criteria.

Representative stable associations included CHIT-1 in lung (Pearson r = 0.887; Spearman ρ = 0.888), TNFRSF1B in transverse colon (r = 0.782; ρ = 0.761), and CHIT-1 in kidney cortex (r = 0.6995; ρ = 0.7270) (Figure 2A–C). The kidney CHIT-1 Pearson coefficient was below 0.70 before rounding and therefore did not meet the stronger both-method criterion.

**Figure 2.**
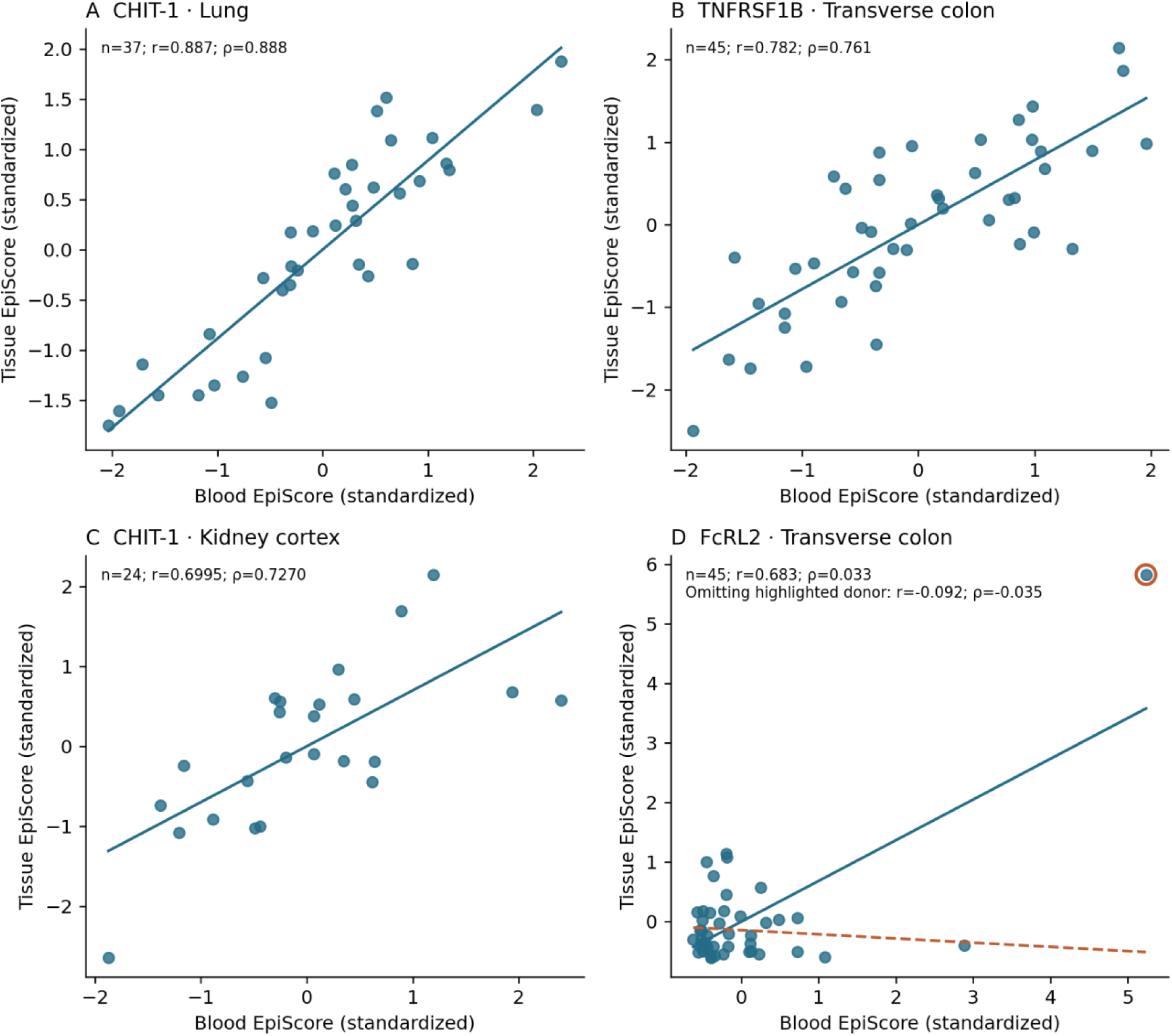
Representative donor-level associations. Paired blood and tissue values are standardized within each comparison. Solid lines show full-sample linear fits. In panel D, the orange ring identifies the donor whose omission produces the largest absolute Pearson change; the dashed line shows the remaining-data fit. No donor was removed from primary analyses. Panel C displays four decimal places to show that the kidney Pearson coefficient is below 0.70 before rounding. In panel D, the same highlighted donor drives the adjusted association: partial r changes from 0.689 to −0.076 when that donor is omitted and the models are refitted.

An additional 18 comparisons met the ≥0.50 threshold for Pearson alone, and 15 met it for Spearman alone. The transverse colon FcRL2 EpiScore illustrated sensitivity to an individual donor: its full-sample Pearson correlation was 0.683, whereas its Spearman correlation was 0.033. Omitting the most influential donor changed these coefficients to −0.092 and −0.035, respectively (Figure 2D). Full-sample estimates were retained as the primary results. The same donor influenced the covariate-adjusted association: partial r was 0.689 in the full sample and −0.076 after omitting that donor and refitting both residualization models.

### Covariate-adjusted analyses retained substantial lung correspondence

Partial Pearson correlations were estimable for 94 scores in each of lung, transverse colon, kidney cortex, and prostate. Lung and colon models included age, sex, BMI, and donor cohort; kidney models included age, sex, and BMI; and prostate models included age, BMI, and donor cohort.

Median absolute partial correlations were 0.522 in lung, 0.255 in transverse colon, 0.263 in kidney cortex, and 0.103 in prostate, compared with unadjusted medians of 0.594, 0.318, 0.286, and 0.092, respectively. After adjustment across the 376 estimable partial-correlation tests, 68 lung, 17 colon, 11 kidney, and one prostate score met global q <0.05. Breast and ovary did not meet the minimum sample-size requirement. These results represent tissue-specific sensitivity analyses because the eligible covariate sets differed across tissues (Supplementary Figure S2). Among initially concordant pairs, partial r ≥0.50 was retained in 48/56 lung, 3/10 colon, 1/2 kidney, and 0/3 prostate comparisons; global partial-correlation q <0.05 was retained in 54/56, 4/10, 1/2, and 0/3, respectively. Both adjusted criteria were retained in 48/56, 3/10, 1/2, and 0/3. Within the initially stable subset, the corresponding counts meeting both adjusted criteria were 42/49 in lung, 2/8 in colon, 0/1 in kidney, and 0/2 in prostate. These are retention summaries of the original concordant and stable subsets, not new leave-one-out tests of the adjusted models.

### Lung concordance extended across secretion categories

The 109 EpiScores were classified by target-protein annotation as secreted to blood (48 scores), not secreted to blood (29), locally or extracellularly secreted but not to blood (15), other or ambiguously secreted (three), or unclear or missing annotation (14). After excluding non-estimable scores, these categories contained 43, 24, 12, two, and 13 scores per tissue, respectively.

In lung, both correlation coefficients were ≥0.50 for 27 of 43 blood-secreted scores (62.8%), eight of 12 locally or extracellularly secreted scores (66.7%), and 10 of 24 scores annotated as not secreted to blood (41.7%). Both scores in the other or ambiguously secreted category and nine of 13 scores with unclear or missing annotations also met this criterion. Within the blood-secreted, locally secreted, and not-secreted-to-blood categories, 26, seven, and seven comparisons, respectively, additionally met the leave-one-out stability criteria (Figure 3A).

**Figure 3.**
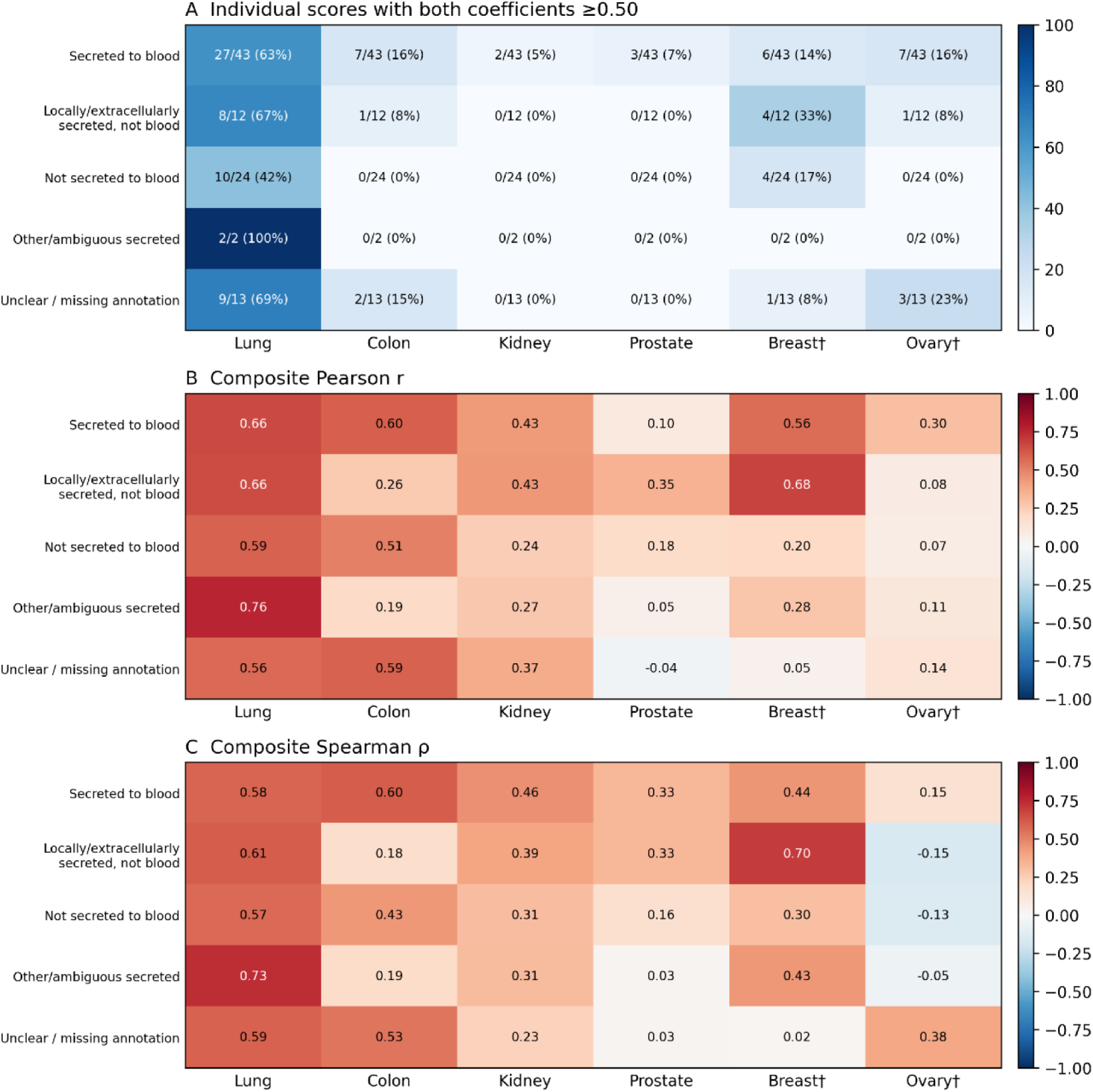
Concordance by target-protein secretion category. A: Individual scores with both Pearson r and Spearman ρ ≥0.50, reported as counts/estimable category totals and percentages. B–C: Signed Pearson and Spearman correlations between donor-level secretion-category composites. The exact 109-score mapping and category results are supplied in Supplementary Data S2. Category comparisons are descriptive. † Exploratory small-sample tissue.

In transverse colon, seven of 43 blood-secreted scores (16.3%) met the both-method concordance criterion, compared with one of 12 locally or extracellularly secreted scores (8.3%) and none of the 24 scores annotated as not secreted to blood. The two qualifying kidney cortex scores and three qualifying prostate scores were all annotated as secreted to blood.

All five lung secretion-category composites had Pearson and Spearman correlations ≥0.50. The blood-secreted composite had coefficients of 0.660 and 0.584, respectively, compared with 0.656 and 0.608 for the locally or extracellularly secreted composite and 0.590 and 0.569 for the not-secreted-to-blood composite. The other or ambiguously secreted composite had the highest coefficients (0.764 and 0.735), although it contained only two estimable scores (Figure 3B–C).

In the transverse colon, the blood-secreted and unclear-annotation composites met the both-method criterion. In breast mammary tissue, only the locally or extracellularly secreted composite met this criterion (Pearson r = 0.678; Spearman ρ = 0.697). No secretion-category composite reached a positive correlation of 0.50 by either method in kidney cortex, prostate, or ovary. Category comparisons were descriptive, with complete estimates provided in Supplementary Data S2.

### Functional-group composites showed heterogeneous correspondence

The 109 EpiScores were assigned to 10 manually curated functional groups. Estimable group sizes ranged from two scores in the coagulation, platelet, and hemostasis group to 20 in the chemokine, cytokine, and growth-factor group.

In lung, individual scores meeting the both-method ≥0.50 criterion were represented in all 10 groups. The chemokine, cytokine, and growth-factor group contained 15 qualifying scores out of 20 (75.0%), while the protease and protease-inhibitor group contained seven out of 10 (70.0%). Fourteen and six of these comparisons, respectively, also met the leave-one-out stability criteria (Figure 4A).

**Figure 4.**
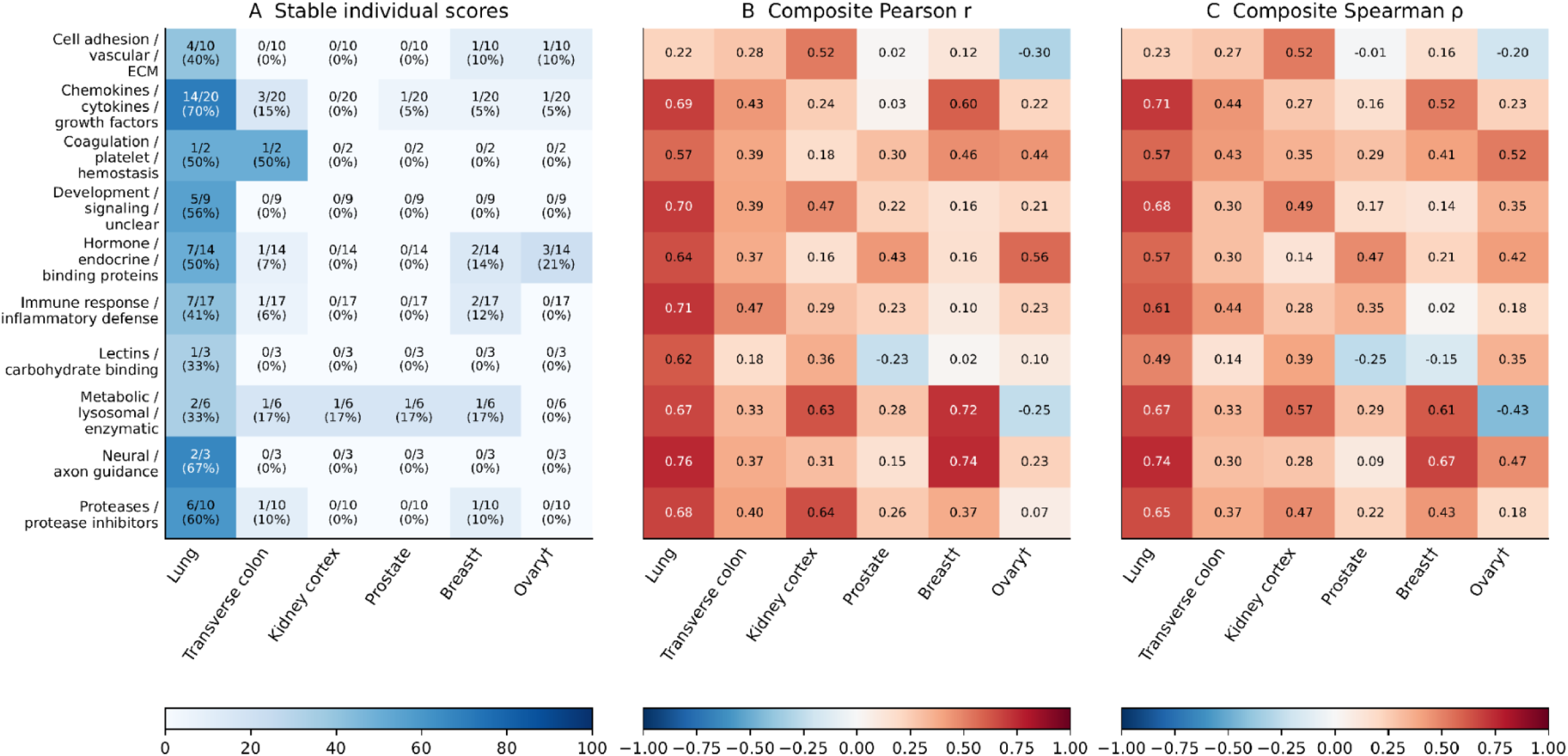
Functional-group concordance. A: Stable individual scores as counts/estimable group totals and percentages. B–C: Signed Pearson and Spearman correlations between donor-level composites. Composite members are standardized separately in matched blood and tissue samples before averaging. These analyses are descriptive, not pathway-enrichment tests. † Exploratory tissue.

Eight of the 10 lung functional-group composites had both coefficients ≥0.50. The neural and axon-guidance composite had the highest Pearson and Spearman correlations (0.755 and 0.743), although it contained only three estimable scores. Other concordant composites included chemokines, cytokines, and growth factors (0.688 and 0.709), metabolic, lysosomal, and enzymatic proteins (0.670 and 0.671), and proteases and protease inhibitors (0.678 and 0.652). The cell-adhesion, vascular, and extracellular-matrix composite had lower correlations (0.221 and 0.234), despite five of its 10 individual scores meeting the concordance criterion and four meeting the stability criteria (Figure 4B–C).

Two kidney cortex composites met the both-method criterion: metabolic, lysosomal, and enzymatic proteins (Pearson r = 0.625; Spearman ρ = 0.566) and cell adhesion, vascular, and extracellular matrix (r = 0.518; ρ = 0.517). Three breast mammary tissue composites met the criterion: neural and axon guidance (r = 0.744; ρ = 0.673), metabolic, lysosomal, and enzymatic proteins (r = 0.723; ρ = 0.612), and chemokines, cytokines, and growth factors (r = 0.595; ρ = 0.515). No functional-group composite met the both-method criterion in transverse colon, prostate, or ovary.

In the descriptive summary across the five tissues with at least 10 matched donors, the metabolic, lysosomal, and enzymatic group had the highest median absolute composite correlations (Pearson, 0.625; Spearman, 0.566). This group also had the highest Pearson correlation in the separate pooled analysis of within-tissue-centered composites (r = 0.439; ρ = 0.435). Pooled estimates were treated as descriptive because the 158 blood–tissue pairs included repeated contributions from the same donors. Complete composite results are provided in Supplementary Data S1.

### Exploratory subgroup analyses did not establish cohort differences

Age-stratified analyses showed relatively high median absolute Pearson correlations in both lung strata: 0.621 among donors aged ≤49 years (n = 21) and 0.541 among those aged >49 years (n = 16). In the transverse colon, the median was 0.391 in the older stratum (>53 years; n = 21) and 0.214 in the younger stratum (≤53 years; n = 24). Kidney cortex and prostate also had higher descriptive medians in their older strata, although all four subgroup medians were below 0.30. Breast and ovary age strata contained only four or five donors each and did not meet the minimum sample size for stratified correlation estimation.

Patterns by donor cohort differed descriptively across tissues. In lung, the median absolute Pearson correlation was 0.731 in organ procurement organization donors (n = 15) and 0.493 in postmortem donors (n = 22). Transverse colon showed the opposite ordering, with medians of 0.241 in organ donors (n = 16) and 0.366 in postmortem donors (n = 29).

Exploratory formal comparisons of signed Pearson correlations were conducted for each of the 94 estimable EpiScores in lung and transverse colon. Ten lung comparisons and two colon comparisons had unadjusted p-values <0.05. None remained significant after Benjamini–Hochberg correction across all 188 tests; the minimum adjusted p-value was 0.172. Thus, no individual EpiScore showed a cohort difference supported after multiple-testing correction.

Prostate median absolute correlations were 0.279 in organ donors and 0.211 in postmortem donors, but this tissue did not meet the eligibility criteria for formal cohort comparisons. All matched kidney cortex and ovary samples belonged to the postmortem cohort. The breast organ-donor stratum contained only two donors.

Sex-stratified analyses were limited by unequal sample sizes. Lung included 31 male donors, with a median absolute Pearson correlation of 0.572, and six female donors. Female strata in transverse colon and kidney cortex each contained eight donors, and breast mammary tissue included five donors of each sex. These smaller strata did not meet the minimum of 10 donors for stratified correlation estimation. Recorded ethnicity strata were also uneven and frequently too small for estimation; these results do not support conclusions about ethnicity-related differences.

All subgroup analyses were exploratory. Formal cohort comparisons were added after inspection of the descriptive results, whereas age, sex, and ethnicity comparisons remained descriptive. The absence of cohort differences meeting the multiple-testing threshold was not interpreted as evidence of equivalent correlations.

## Discussion

This study evaluated whether blood-trained protein EpiScores preserve between-donor variation when applied to DNAm profiles from six non-blood tissues. Lung showed the broadest observed blood–tissue correspondence, including the largest number of comparisons meeting the positive-concordance and leave-one-out stability criteria. Other tissues showed more selective patterns. Across all tissues, 97 protein–tissue comparisons met the both-method concordance criterion, and 73 additionally met the operational stability criteria. These findings support evaluating cross-tissue correspondence for individual EpiScores and tissue combinations rather than assuming that the entire panel behaves similarly across tissues. Because donor sets and sample sizes differed, the observed tissue ordering should be interpreted descriptively.

Previous studies have primarily evaluated protein EpiScores through their associations with circulating protein measurements and health outcomes [4,5,7,8]. Our analysis addresses a complementary question: whether applying the same prediction model to blood and another tissue yields corresponding variation among matched donors. This distinction matters for biomarker interpretation. A blood EpiScore could be associated with disease through systemic influences without closely corresponding to the same score in the affected tissue. Conversely, strong blood–tissue correspondence does not establish that either score accurately measures local protein abundance. It identifies shared variation in a methylation-derived signature whose biological and clinical meaning requires further investigation.

The importance of biological context is consistent with previous evaluations of protein EpiScores across the lifecourse and DNAm biomarkers across tissues [9,10,14]. These studies motivate testing whether properties established in one setting persist in another. Our donor-matched analysis extends this question to multiple internal tissues and identifies specific score–tissue combinations for subsequent validation.

### Interpreting the broad lung correspondence

The lung pattern was evident across individual EpiScores, leave-one-out analyses, and several functional-group composites. Lung also retained substantial correspondence after adjustment for available covariates, with 48 of 56 initially concordant comparisons retaining partial r ≥0.50 and global q <0.05. Most initially concordant comparisons in colon, kidney cortex, and prostate did not retain both adjusted criteria; breast and ovary could not be evaluated because of sample size. These results prioritize lung for follow-up while leaving the sources of blood–lung covariance unresolved.

Several explanations are plausible. Methylation in blood and lung may reflect shared exposures, donor characteristics, or immune-related variation. Correlated differences in cellular composition could contribute if immune-cell abundance varies across compartments. Shared genetic influences on methylation provide another possible explanation, given the cross-tissue methylation-QTL architecture investigated in GTEx [11]. These mechanisms may operate simultaneously. Distinguishing them will require analyses of the CpGs contributing to individual scores, their genetic associations, and their variation across cell populations. Blood retained within lung tissue and variation in resident or infiltrating immune cells could contribute to these relationships. Without cell-type deconvolution, correspondence cannot be attributed specifically to lung parenchymal cells.

### Biological relevance of selected score–tissue findings

The biological functions of selected target proteins provide context for prioritizing the observed score– tissue relationships. In lung, concordant EpiScores included proteins involved in airway host defense and epithelial function. CHIT-1 is the principal active chitinase in the human lung, with activity influenced by genotype and smoking [15]. The polymeric immunoglobulin receptor (pIgR) participates in mucosal antibody transport, and studies of chronic obstructive pulmonary disease have linked its reduced epithelial expression to impaired airway immune defense [16]. The strong, statistically significant, and leave-one-out-stable correspondence of these scores motivates investigation of whether shared immune-related influences contribute to their methylation signatures across blood and lung.

Other concordant lung targets connect to inflammatory signaling and tissue remodeling. CCL17 and CCL22 have been investigated in idiopathic pulmonary fibrosis, including their relationship to CCR4-expressing inflammatory cells [17]. Experimental work also implicates the Sema3E–Plexin D1 axis in fibroblast activation and lung fibrosis [18]. These findings suggest specific processes to examine in follow-up studies of the corresponding EpiScores. However, the present analysis did not test pulmonary disease mechanisms, and the broad lung pattern could reflect several shared influences rather than one common pathway.

In the transverse colon, TNFRSF1B was a particularly consistent candidate, meeting the positive-concordance, global FDR, and leave-one-out stability criteria for both correlation methods. TNFRSF1B encodes TNFR2, which has an experimentally demonstrated role in intestinal epithelial recovery: a study of colitis and epithelial models found that TNFR2 helps terminate regenerative signaling and restore epithelial homeostasis [19]. This provides a biological rationale for investigating the score in relation to epithelial repair and inflammatory context. It does not establish that the blood–colon score association measures TNFR2 activity or that the donors had intestinal inflammation.

In the kidney cortex, CHIT-1 met both-method significance and leave-one-out stability criteria. A study of patients with type 2 diabetes reported associations of chitotriosidase enzyme activity and genotype with nephropathy, linking the measured protein to a renal disease setting [20]. Adiponectin provided an additional, less consistent candidate: both correlations reached the positive-concordance threshold, but only Pearson passed global FDR correction, and the pair did not meet the leave-one-out stability criteria. Experimental kidney-specific gain-and loss-of-function studies in mice have shown that locally produced adiponectin influences fatty acid utilization and renal gluconeogenesis [21]. Together, these examples motivate investigation of immune and metabolic influences on renal EpiScores, while retaining the distinction between the more consistent CHIT-1 finding and the exploratory adiponectin result.

In the prostate, CHIT-1 and CCL18 met the both-method significance and stability criteria, despite the generally limited correspondence across the panel. Previous clinical research examined serum chitotriosidase activity in primary prostate cancer and benign prostatic hyperplasia [22], providing a direct connection between the CHIT-1 target and prostate-related biomarker research. Its recurrence across several tissues in the present study also raises a broader possibility: the EpiScore may capture shared donor-level influences rather than a prostate-specific process. Distinguishing those possibilities would require direct protein measurements, assessment of cellular composition, and evaluation in clinically characterized samples.

In breast, STC1 illustrates why statistical significance, correlation magnitude, and stability should be reported separately. Its EpiScore showed positive blood–breast correspondence (Pearson r = 0.646; Spearman ρ = 0.697) and met the leave-one-out stability criteria, although neither correlation passed global FDR correction. Experimental studies in breast cancer cells have linked STC1 to homologous recombination-mediated DNA repair and resistance to DNA-damaging treatments [23]. This literature supports retaining STC1 as a biologically motivated exploratory candidate, without presenting it as a statistically confirmed association or evidence that the score reflects malignant processes in GTEx breast tissue. Replication is particularly important given the ten matched donors.

In ovary, SHBG showed positive correspondence (Pearson r = 0.721; Spearman ρ = 0.800) and met the leave-one-out stability criteria, with global FDR significance for Spearman only. SHBG has been detected and characterized in human follicular fluid as well as serum, providing a direct link to the ovarian steroid environment [24]. This supports follow-up of the score in an endocrine context, but does not establish local synthesis, ovarian hormone production, or a specific reproductive mechanism. With only nine matched donors, the present result remains a candidate for replication rather than a validated ovarian biomarker.

These examples connect the observed patterns to pulmonary host defense, intestinal repair, renal metabolism, and reproductive biology. They provide a rationale for selecting follow-up measurements, rather than an independent test of the EpiScores. The cited studies concern target proteins, enzyme activities, or experimentally manipulated pathways; they do not validate the corresponding DNAm signatures as measures of those processes. Matched methylation and protein measurements will be needed to determine whether the observed correspondence reflects protein abundance, related biological variation, or shared upstream influences.

### Secretion annotations and composite scores

Lung concordance extended across proteins annotated as secreted to blood, locally or extracellularly secreted, and not secreted to blood. Thus, observed correspondence was not confined to the blood-secreted category. However, these descriptive comparisons do not establish whether secretion status predicts concordance or accounts for differences between categories. Protein localization, circulating abundance, and the methylation features comprising an EpiScore are distinct properties. Their relationships require direct investigation rather than inference from annotation alone.

Functional-group composites also showed that individual-score correspondence and aggregate correspondence need not coincide. Eight lung composites met the both-method concordance criterion, yet the cell-adhesion, vascular, and extracellular-matrix composite had relatively weak correlations despite several concordant individual members. Kidney cortex showed the converse pattern, with selected concordant composites despite few concordant individual scores.

These patterns are compatible with the construction of the composites: their correlations depend on covariance among all contributing scores, including cross-tissue relationships between different group members. Averaging may emphasize shared variation or dilute signals confined to particular members. Individual and composite analyses therefore describe different aspects of the data. The manually curated groups provide exploratory summaries and should not be interpreted as evidence of pathway enrichment or validated measures of pathway activity.

### Sensitivity to individual donors and covariates

Combining Pearson and Spearman correlations with leave-one-out analysis helped distinguish consistent sample patterns from associations sensitive to individual observations. Approximately three quarters of initially concordant comparisons met the stability criteria, while one quarter did not. The transverse colon FcRL2 example illustrates the importance of these checks: a relatively high Pearson correlation coexisted with a near-zero Spearman correlation, and the Pearson association disappeared after the omission of one influential donor.

These findings support reporting both correlation methods and examining donor influence when prioritizing EpiScores in modest samples. Nevertheless, the thresholds used here are operational criteria, and stability under single-donor omission is not equivalent to independent replication. Likewise, statistical significance, correlation magnitude, and leave-one-out stability describe different properties and should not be treated interchangeably.

Covariate-adjusted results provide additional sensitivity evidence but do not isolate a biological mechanism. Eligible covariate sets differed across tissues, and adjustment could not address all potential sources of shared variation. Exploratory organ-donor versus postmortem comparisons did not survive multiple-testing correction. This provides insufficient evidence for individual-score cohort differences under the specified testing framework, rather than evidence that the cohorts have equivalent correlations. The descriptive lung cohort contrast was substantial (median |r| 0.731 in organ procurement organization donors versus 0.493 in postmortem donors), but the groups differed in size and no individual-score contrast passed FDR correction. Collection context, ischemic time, agonal circumstances, and immune-cell content are plausible contributors that were not separated by the present analysis.

### Strengths and limitations

A principal strength is the use of donor-matched blood and tissue samples, allowing direct assessment of whether EpiScores preserve relative differences among the same individuals. The analysis also distinguishes positive correlation magnitude from statistical significance, evaluates sensitivity to individual donors, and provides complete tissue-specific results alongside significant-only lists. The documented downstream workflow supports verification and regeneration of the reported analyses from the score exports and metadata.

Several limitations constrain interpretation. First, we compared methylation-derived scores rather than measured proteins. Correlation describes between-donor correspondence and does not establish agreement in absolute score values, calibration, or prediction of tissue protein abundance. Second, matched samples were unequal and sometimes small. Sampling variability limits the precision of individual correlations and complicates comparisons of absolute-correlation summaries across tissues, particularly for breast and ovary. No formal analysis established differences between tissue-wide correlation distributions. The analytical sample was predominantly male (45/54 donors), and five of the ten breast donors were male, limiting generalization to female breast biology. The source GTEx methylation cohort was predominantly of European ancestry [11]; genetic ancestry was not independently inferred for our matched subset.

Third, some donors contributed to multiple tissue comparisons. Pooled composite results were therefore treated as descriptive, without inference that assumed independent observations. Partial correlations addressed selected donor covariates but not cellular composition or genetic effects, and residual confounding remains possible. The 109 predictors also represent a selected published panel; these findings should not be generalized to all protein-associated DNAm signatures. No cell-type deconvolution was performed, and ischemic time and agonal factors were not included in the adjustment models. Blood or immune-cell content may therefore contribute to the lung pattern. Low-variance scores may be sensitive to measurement noise, while highly correlated or near-duplicate scores could contribute redundant information to panel summaries and composites; the analysis did not formally quantify effective independence among predictors.

Predictor CpG coverage is an additional limitation. Although full processed matrices containing 754,119 CpG sites were submitted for each tissue, availability was not quantified separately for each protein model. MethylDetectR’s substitution of reference means for absent predictor CpGs permits score calculation but does not recover donor-specific information at those sites. An additive constant does not itself induce correlation, but omission of informative CpG variation may affect the resulting score relationships. We cannot determine whether this contributed to the 15 non-estimable EpiScores. A model-specific audit of available CpGs and prediction weights would help clarify this issue. The hosted application’s version identifier was not recorded, and the reproducible downstream workflow does not automatically replay the original scoring step.

Finally, the biological annotations and targeted literature review were assembled after examination of the findings. They provide hypotheses rather than independent validation, and the literature review was not systematic. Functional-group assignments and composites likewise require evaluation in independent data before being interpreted as biological biomarkers.

### Future directions

Priorities include replication in independent donor-matched samples, measurement of the corresponding proteins in both blood and tissue, and audits of score-specific CpG availability and prediction weights. Larger samples would permit more precise assessment of tissue differences and the influence of collection context and donor characteristics. Integrating cellular composition, methylation-associated genetic variation, and environmental exposures could help distinguish sources of cross-tissue covariance. For replicated score–tissue combinations, prospective studies should evaluate whether EpiScores improve prediction of tissue-relevant phenotypes or health outcomes beyond established predictors.

## Conclusion

Blood-trained protein EpiScores showed selective cross-tissue correspondence in matched GTEx donors. Of 564 estimable protein–tissue comparisons, 97 met the both-method positive-concordance criterion, and 73 of these also met the leave-one-out stability criteria. Lung showed the broadest observed correspondence, with substantial correlations retained after adjustment for available donor covariates. These findings prioritize specific score–tissue combinations for independent replication and validation against measured proteins. They do not establish that blood EpiScores measure local tissue protein abundance. Outside lung, most initially concordant comparisons in tissues eligible for adjustment did not retain both partial r ≥0.50 and q <0.05; the small breast and ovary samples precluded evaluation of covariate sensitivity.

## Declarations

### Ethics and consent

This study used publicly available, de-identified processed GTEx methylation data from GEO accession GSE213478 [11]. No new participants were recruited or specimens collected. No controlled -access data-use agreement or additional institutional review board approval was required for this secondary analysis.

### Data and code availability

Processed DNA methylation data are publicly available through GEO accession GSE213478. Aggregate results and annotation mappings are provided in Supplementary Data S1 and S2. Analysis scripts, publication-figure code, and derived analysis data are available from the first author upon reasonable request.

### Author Contributions

Paulina Gonzalez Orozco conducted the analyses, generated the figures and tables, and drafted the manuscript. Fredrick Schumacher supervised the research and critically revised the manuscript. Both authors contributed to study design and interpretation of the results.

## Supplementary materials

Supplementary Figures S1–S2 show Spearman correlations and partial-correlation sensitivity results. Supplementary Tables S1–S6 provide tissue-specific lists of proteins significant in at least one correlation method after global Benjamini–Hochberg adjustment. Supplementary Data S1 contains the complete 109-score results for each tissue, including non-estimable comparisons. Supplementary Data S2 provides the secretion-category map, category summaries, composite results, and numerical reconciliation.

**Supplementary Figure S1.**
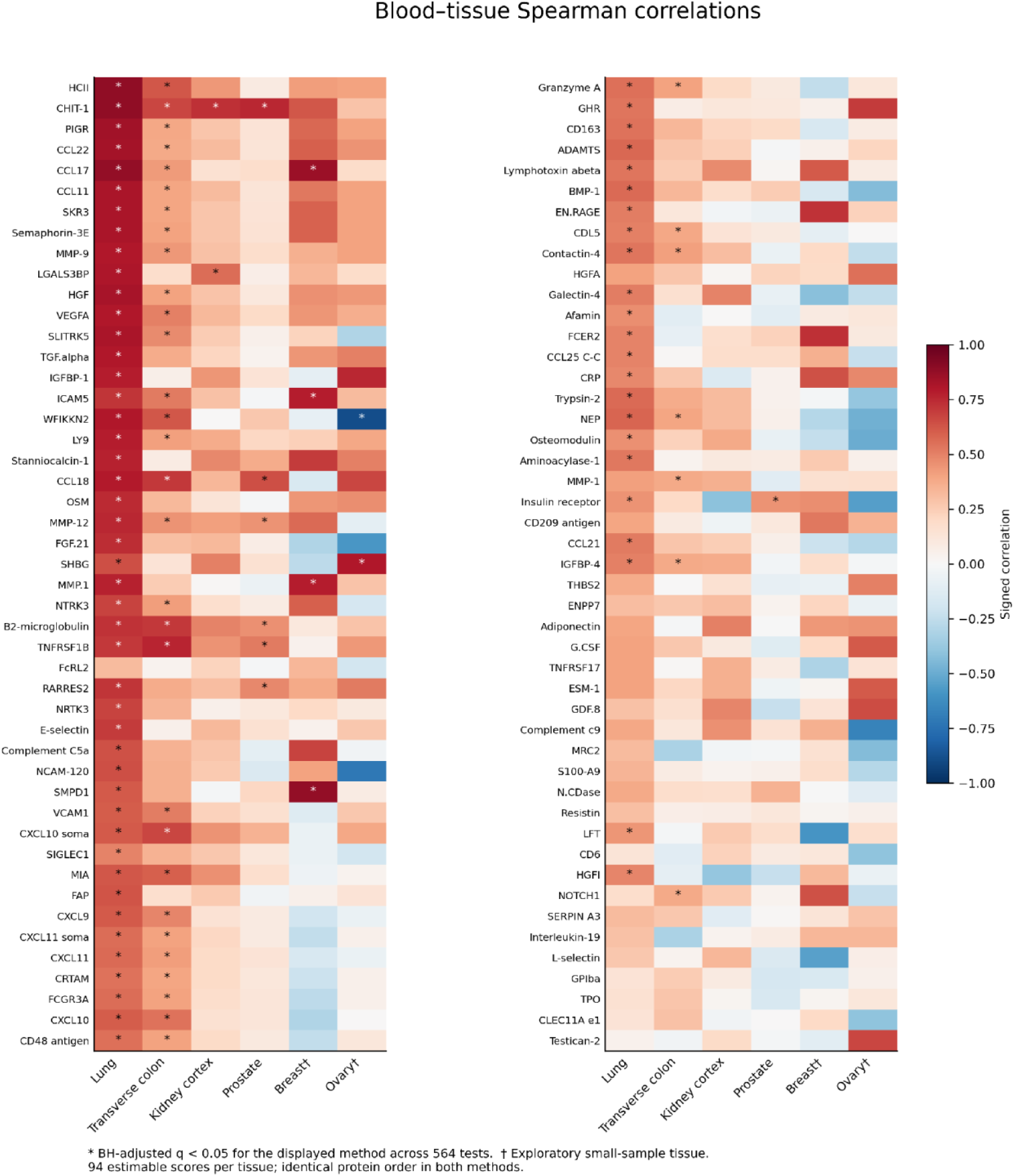
Spearman correlations. Signed Spearman correlations in the same protein order as Figure 1A. Asterisks indicate global BH-adjusted q <0.05 for Spearman across 564 tests. † Exploratory tissue.

**Supplementary Figure S2.**
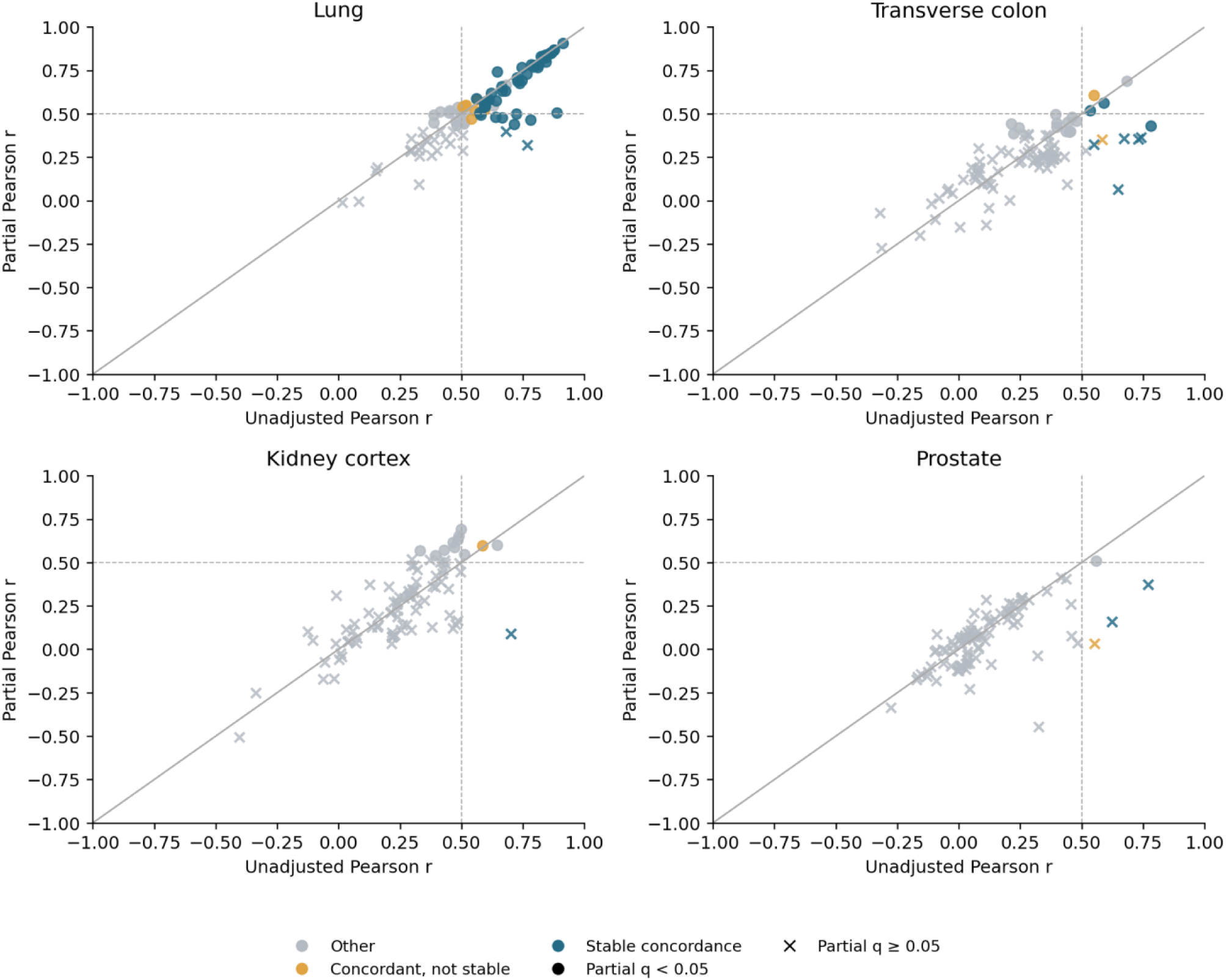
Covariate-adjusted sensitivity. Each point represents one of 94 estimable protein EpiScores per tissue. Signed full-sample Pearson r is plotted against partial Pearson r. Colors distinguish scores meeting the original both-method concordance and leave-one-out stability criteria; circles mark global partial-correlation q <0.05 and crosses mark q ≥0.05. The diagonal denotes equal adjusted and unadjusted coefficients; dashed lines mark r = 0.50. Covariate sets differ by tissue as described in Methods. Breast and ovary were ineligible for adjusted analysis. Classification colors refer to the unadjusted analyses and do not imply stability of adjusted models.

### Significant blood–tissue protein EpiScore correlations

#### Supplementary Tables S1–S6

These tables list every protein EpiScore significant in at least one method (global Benjamini–Hochberg q <0.05). Adjustment was performed separately for Pearson and Spearman across all 564 estimable protein–tissue comparisons before selecting these rows. Signed coefficients are retained, including inverse associations. Statistical significance is distinct from the positive-concordance magnitude thresholds and from leave-one-out stability.

Protein labels are the original MethylDetectR score identifiers. Pearson confidence intervals are nominal 95% Fisher-z intervals, not simultaneous or multiplicity-adjusted intervals. “Both” denotes significance in both methods; “P only” and “S only” denote Pearson-only and Spearman-only significance. “Stable” requires both coefficients ≥0.50 in the full sample and after every donor omission, with maximum absolute change <0.30 for each coefficient. It does not denote independent replication. Values are rounded for display only. Partial r and partial q are from the covariate-adjusted sensitivity analyses; partial q uses global BH correction across 376 tests, independently of row selection. A dash indicates that adjusted analysis was ineligible. Kidney CHIT-1 coefficients are displayed to four decimal places to avoid rounding its Pearson coefficient above the strong-concordance threshold.

#### Included results

- Lung: 74 proteins; 67 significant in both methods, 5 in Pearson only, and 2 in Spearman only.
- Transverse colon: 42 proteins; 26 significant in both methods, 5 in Pearson only, and 11 in Spearman only.
- Kidney cortex: 4 proteins; 1 significant in both methods, 2 in Pearson only, and 1 in Spearman only.
- Prostate: 8 proteins; 6 significant in both methods, 1 in Pearson only, and 1 in Spearman only.
- Breast mammary tissue: 4 proteins; 1 significant in both methods, 0 in Pearson only, and 3 in Spearman only.
- Ovary: 2 proteins; 0 significant in both methods, 0 in Pearson only, and 2 in Spearman only.

**Table S1.**
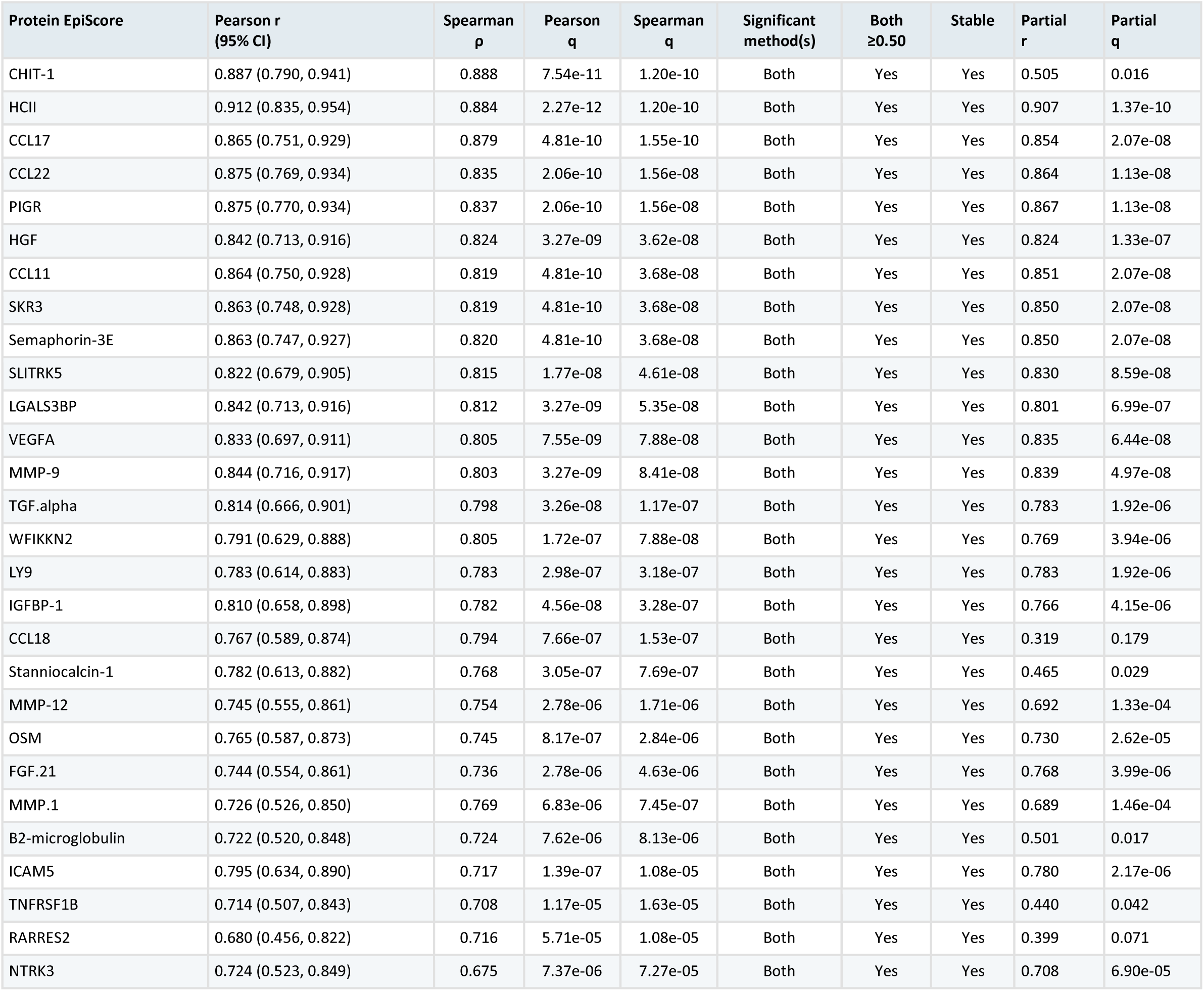
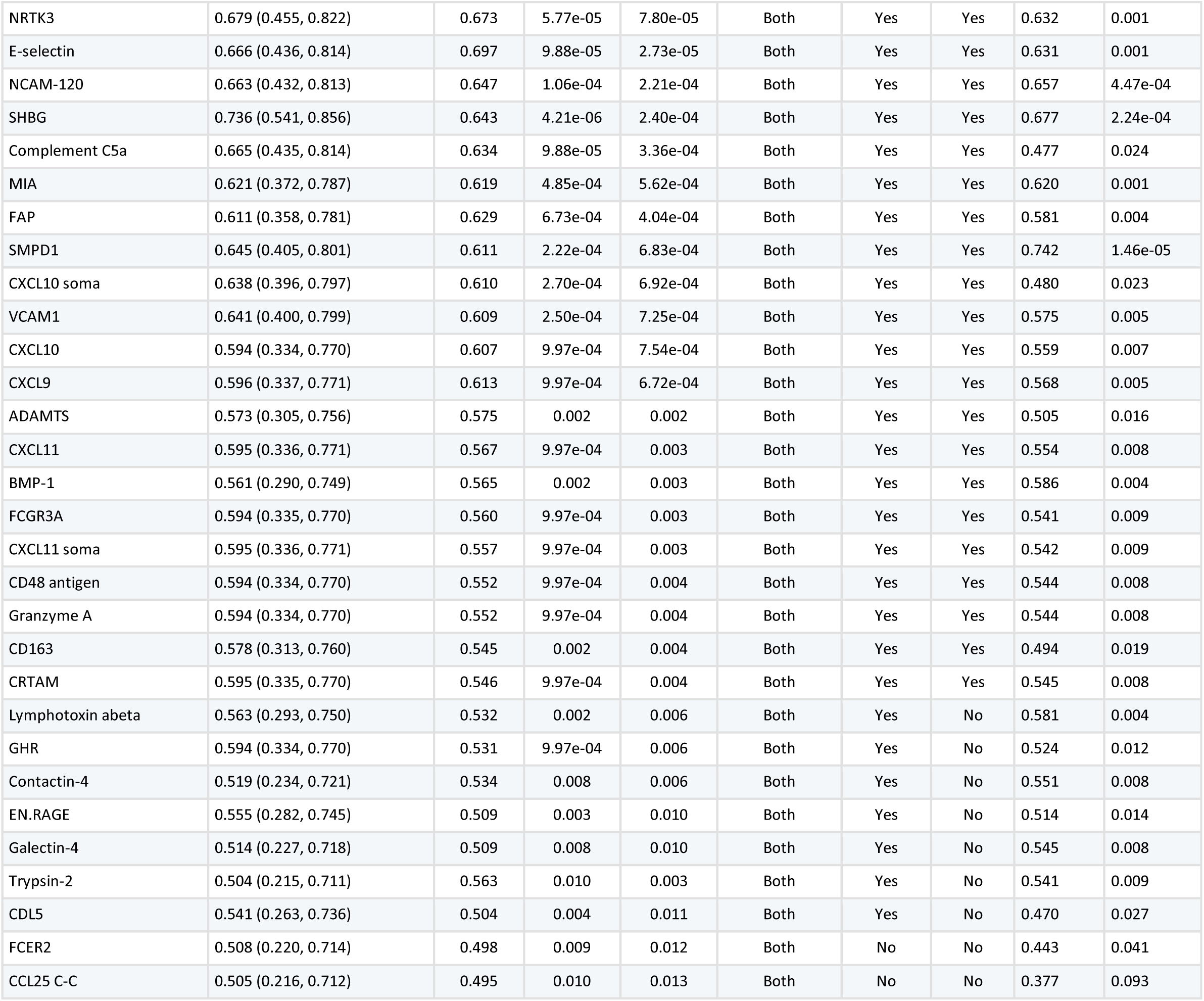
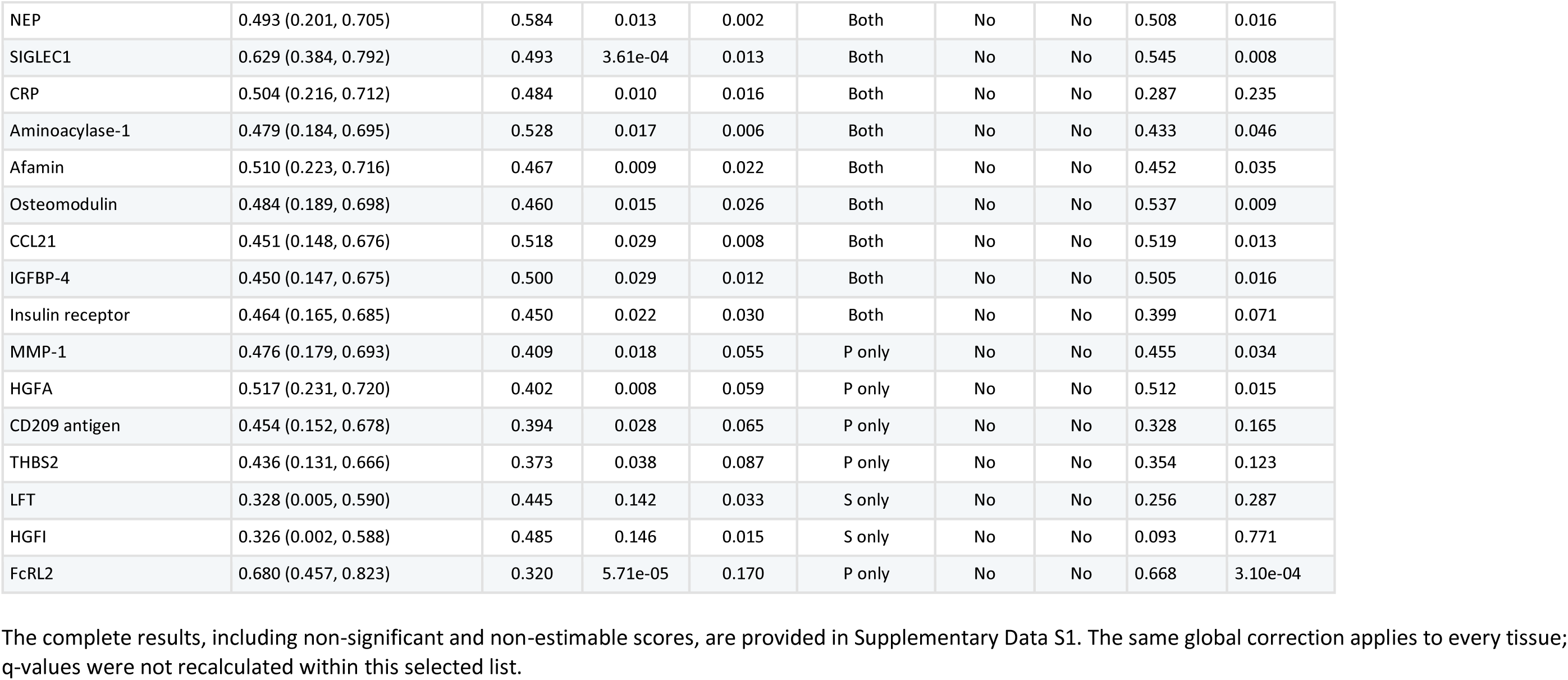
Lung. 37 matched donors; 74 of 94 estimable scores significant in at least one method.

**Table S2.** Transverse colon. 45 matched donors; 42 of 94 estimable scores significant in at least one method.

| Protein EpiScore | Pearson r<br>(95% CI) | Spearman<br>ρ | Pearson<br>q | Spearman<br>q | Significant<br>method(s) | Both<br>≥0.50 | Stable | Partial<br>r | Partial<br>q |
| --- | --- | --- | --- | --- | --- | --- | --- | --- | --- |
| TNFRSF1B | 0.782 (0.634, 0.875) | 0.761 | 9.79e-09 | 5.98e-08 | Both | Yes | Yes | 0.431 | 0.024 |
| B2-microglobulin | 0.740 (0.571, 0.849) | 0.707 | 1.85e-07 | 1.37e-06 | Both | Yes | Yes | 0.365 | 0.066 |
| CCL18 | 0.729 (0.554, 0.842) | 0.707 | 3.49e-07 | 1.37e-06 | Both | Yes | Yes | 0.354 | 0.073 |
| CXCL10 soma | 0.672 (0.471, 0.806) | 0.670 | 7.62e-06 | 9.50e-06 | Both | Yes | Yes | 0.356 | 0.073 |
| CHIT-1 | 0.648 (0.438, 0.791) | 0.667 | 2.39e-05 | 1.08e-05 | Both | Yes | Yes | 0.065 | 0.806 |
| HCII | 0.590 (0.359, 0.753) | 0.620 | 2.56e-04 | 8.36e-05 | Both | Yes | Yes | 0.562 | 0.002 |
| MIA | 0.549 (0.305, 0.726) | 0.596 | 9.97e-04 | 2.21e-04 | Both | Yes | No | 0.608 | 3.51e-04 |
| CXCL10 | 0.550 (0.306, 0.727) | 0.541 | 9.97e-04 | 0.001 | Both | Yes | Yes | 0.322 | 0.118 |
| WFIKKN2 | 0.534 (0.285, 0.715) | 0.623 | 0.001 | 7.52e-05 | Both | Yes | Yes | 0.519 | 0.005 |
| VCAM1 | 0.583 (0.350, 0.749) | 0.514 | 3.21e-04 | 0.003 | Both | Yes | No | 0.353 | 0.074 |
| CXCL9 | 0.517 (0.263, 0.704) | 0.486 | 0.002 | 0.006 | Both | No | No | 0.289 | 0.174 |
| SLITRK5 | 0.461 (0.194, 0.665) | 0.475 | 0.010 | 0.008 | Both | No | No | 0.480 | 0.010 |
| VEGFA | 0.454 (0.186, 0.660) | 0.504 | 0.011 | 0.004 | Both | No | No | 0.399 | 0.041 |
| MMP-12 | 0.449 (0.179, 0.656) | 0.451 | 0.013 | 0.013 | Both | No | No | 0.395 | 0.042 |
| ICAM5 | 0.443 (0.171, 0.652) | 0.516 | 0.015 | 0.003 | Both | No | No | 0.453 | 0.017 |
| MMP-9 | 0.436 (0.163, 0.647) | 0.436 | 0.017 | 0.017 | Both | No | No | 0.398 | 0.041 |
| CXCL11 | 0.429 (0.155, 0.642) | 0.421 | 0.019 | 0.025 | Both | No | No | 0.253 | 0.240 |
| CXCL11 soma | 0.399 (0.119, 0.620) | 0.403 | 0.037 | 0.033 | Both | No | No | 0.250 | 0.243 |
| NTRK3 | 0.397 (0.118, 0.619) | 0.416 | 0.038 | 0.027 | Both | No | No | 0.423 | 0.027 |
| CCL17 | 0.388 (0.106, 0.612) | 0.428 | 0.044 | 0.021 | Both | No | No | 0.320 | 0.118 |
| FCGR3A | 0.388 (0.106, 0.612) | 0.404 | 0.044 | 0.033 | Both | No | No | 0.242 | 0.264 |
| CD48 antigen | 0.386 (0.104, 0.610) | 0.404 | 0.045 | 0.033 | Both | No | No | 0.239 | 0.264 |
| Granzyme A | 0.386 (0.104, 0.610) | 0.404 | 0.045 | 0.033 | Both | No | No | 0.239 | 0.264 |
| CDL5 | 0.385 (0.103, 0.610) | 0.386 | 0.045 | 0.044 | Both | No | No | 0.259 | 0.235 |
| CRTAM | 0.382 (0.100, 0.608) | 0.397 | 0.047 | 0.037 | Both | No | No | 0.241 | 0.264 |
| MMP-1 | 0.394 (0.114, 0.616) | 0.378 | 0.039 | 0.049 | Both | No | No | 0.495 | 0.008 |
| LY9 | 0.373 (0.089, 0.601) | 0.412 | 0.053 | 0.029 | S only | No | No | 0.219 | 0.305 |
| RARRES2 | 0.440 (0.168, 0.650) | 0.369 | 0.016 | 0.057 | P only | No | No | 0.094 | 0.737 |

| Protein EpiScore | Pearson r<br>(95% CI) | Spearman<br>$\rho$ | Pearson<br>q | Spearman<br>q | Significant<br>method(s) | Both<br>$\geq 0.50$ | Stable | Partial<br>r | Partial<br>q |
| --- | --- | --- | --- | --- | --- | --- | --- | --- | --- |
| NOTCH1 | 0.366 (0.081, 0.595) | 0.378 | 0.059 | 0.049 | S only | No | No | 0.335 | 0.097 |
| HGF | 0.365 (0.080, 0.595) | 0.402 | 0.059 | 0.034 | S only | No | No | 0.289 | 0.174 |
| SMPD1 | 0.399 (0.119, 0.620) | 0.355 | 0.037 | 0.067 | P only | No | No | 0.446 | 0.019 |
| Contactin-4 | 0.353 (0.067, 0.586) | 0.409 | 0.068 | 0.030 | S only | No | No | 0.376 | 0.057 |
| PIGR | 0.350 (0.063, 0.583) | 0.383 | 0.071 | 0.046 | S only | No | No | 0.257 | 0.235 |
| CCL11 | 0.342 (0.054, 0.578) | 0.396 | 0.079 | 0.038 | S only | No | No | 0.268 | 0.215 |
| SKR3 | 0.342 (0.054, 0.577) | 0.391 | 0.079 | 0.040 | S only | No | No | 0.258 | 0.235 |
| Semaphorin-3E | 0.341 (0.053, 0.577) | 0.400 | 0.079 | 0.035 | S only | No | No | 0.257 | 0.235 |
| NRTK3 | 0.388 (0.107, 0.612) | 0.341 | 0.044 | 0.083 | P only | No | No | 0.280 | 0.192 |
| CCL22 | 0.312 (0.020, 0.554) | 0.381 | 0.122 | 0.047 | S only | No | No | 0.238 | 0.265 |
| NEP | 0.281 (-0.013, 0.531) | 0.392 | 0.172 | 0.039 | S only | No | No | 0.383 | 0.050 |
| HGFA | 0.478 (0.214, 0.676) | 0.279 | 0.007 | 0.191 | P only | No | No | 0.460 | 0.015 |
| IGFBP-4 | 0.268 (-0.028, 0.520) | 0.379 | 0.201 | 0.049 | S only | No | No | 0.362 | 0.069 |
| FcRL2 | 0.683 (0.487, 0.813) | 0.033 | 4.56e-06 | 0.933 | P only | No | No | 0.689 | 1.28e-05 |

The complete results, including non-significant and non-estimable scores, are provided in Supplementary Data S1. The same global correction applies to every tissue; q-values were not recalculated within this selected list.

**Table S3.** Kidney cortex. 24 matched donors; 4 of 94 estimable scores significant in at least one method.

| Protein EpiScore | Pearson r<br>(95% CI) | Spearman<br>$\rho$ | Pearson<br>q | Spearman<br>q | Significant<br>method(s) | Both<br>$\geq 0.50$ | Stable | Partial<br>r | Partial<br>q |
| --- | --- | --- | --- | --- | --- | --- | --- | --- | --- |
| CHIT-1 | 0.6995 (0.413, 0.860) | 0.7270 | 0.001 | 6.83e-04 | Both | Yes | Yes | 0.089 | 0.815 |
| Adiponectin | 0.584 (0.237, 0.799) | 0.501 | 0.017 | 0.057 | P only | Yes | No | 0.597 | 0.022 |
| Stanniocalcin-1 | 0.645 (0.327, 0.832) | 0.478 | 0.005 | 0.071 | P only | No | No | 0.602 | 0.020 |
| LGALS3BP | 0.424 (0.025, 0.707) | 0.558 | 0.124 | 0.027 | S only | No | No | 0.458 | 0.109 |

The complete results, including non-significant and non-estimable scores, are provided in Supplementary Data S1. The same global correction applies to every tissue; q-values were not recalculated within this selected list.

**Table S4.** Prostate. 33 matched donors; 8 of 94 estimable scores significant in at least one method.

| Protein EpiScore | Pearson r<br>(95% CI) | Spearman<br>$\rho$ | Pearson<br>q | Spearman<br>q | Significant<br>method(s) | Both<br>$\geq 0.50$ | Stable | Partial<br>r | Partial<br>q |
| --- | --- | --- | --- | --- | --- | --- | --- | --- | --- |
| CHIT-1 | 0.769 (0.579, 0.880) | 0.755 | 3.53e-06 | 8.13e-06 | Both | Yes | Yes | 0.375 | 0.118 |
| CCL18 | 0.623 (0.356, 0.796) | 0.647 | 9.97e-04 | 5.98e-04 | Both | Yes | Yes | 0.160 | 0.589 |
| TNFRSF1B | 0.552 (0.258, 0.753) | 0.520 | 0.007 | 0.013 | Both | Yes | No | 0.031 | 0.913 |
| B2-microglobulin | 0.483 (0.167, 0.709) | 0.456 | 0.026 | 0.039 | Both | No | No | 0.035 | 0.906 |
| RARRES2 | 0.455 (0.133, 0.691) | 0.479 | 0.041 | 0.028 | Both | No | No | 0.260 | 0.305 |
| MMP-12 | 0.459 (0.137, 0.693) | 0.449 | 0.039 | 0.043 | Both | No | No | 0.078 | 0.802 |
| Insulin receptor | 0.437 (0.110, 0.678) | 0.450 | 0.052 | 0.043 | S only | No | No | 0.405 | 0.082 |
| HGFA | 0.558 (0.266, 0.757) | 0.224 | 0.006 | 0.452 | P only | No | No | 0.510 | 0.020 |

The complete results, including non-significant and non-estimable scores, are provided in Supplementary Data S1. The same global correction applies to every tissue; q-values were not recalculated within this selected list.

**Table S5.** Breast mammary tissue. 10 matched donors; 4 of 94 estimable scores significant in at least one method. Exploratory small-sample analysis.

| Protein EpiScore | Pearson r<br>(95% CI) | Spearman<br>$\rho$ | Pearson<br>q | Spearman<br>q | Significant<br>method(s) | Both<br>$\geq 0.50$ | Stable | Partial<br>r | Partial<br>q |
| --- | --- | --- | --- | --- | --- | --- | --- | --- | --- |
| SMPD1 | 0.868 (0.525, 0.968) | 0.879 | 0.008 | 0.006 | Both | Yes | Yes | — | — |
| ICAM5 | 0.760 (0.250, 0.940) | 0.782 | 0.051 | 0.039 | S only | Yes | Yes | — | — |
| CCL17 | 0.750 (0.228, 0.937) | 0.855 | 0.056 | 0.012 | S only | Yes | Yes | — | — |
| MMP.1 | 0.624 (-0.009, 0.900) | 0.782 | 0.157 | 0.039 | S only | Yes | Yes | — | — |

The complete results, including non-significant and non-estimable scores, are provided in Supplementary Data S1. The same global correction applies to every tissue; q-values were not recalculated within this selected list.

**Table S6.** Ovary. 9 matched donors; 2 of 94 estimable scores significant in at least one method. Exploratory small-sample analysis.

| Protein EpiScore | Pearson r<br>(95% CI) | Spearman<br>$\rho$ | Pearson<br>q | Spearman<br>q | Significant<br>method(s) | Both<br>$\geq 0.50$ | Stable | Partial<br>r | Partial<br>q |
| --- | --- | --- | --- | --- | --- | --- | --- | --- | --- |
| WFIKKN2 | -0.786 (-0.953, -0.256) | -0.883 | 0.054 | 0.012 | S only | No | No | — | — |
| SHBG | 0.721 (0.110, 0.937) | 0.800 | 0.100 | 0.046 | S only | Yes | Yes | — | — |

The complete results, including non-significant and non-estimable scores, are provided in Supplementary Data S1. The same global correction applies to every tissue; q-values were not recalculated within this selected list.

